# Generation of a *Nr2f2-driven* inducible Cre mouse to target interstitial cells in the reproductive system

**DOI:** 10.1101/2025.01.22.634276

**Authors:** Paula R. Brown, Martin A. Estermann, Artiom Gruzdev, Gregory J. Scott, Thomas B. Hagler, Manas K. Ray, Humphrey Hung-Chang Yao

## Abstract

The orphan nuclear receptor *Nr2f2*, also known as COUP-TFII, play important roles in development and function of multiple organs, including the reproductive system. NR2F2 is expressed in the interstitial cells of both embryonic and adult testes, ovaries, and reproductive tracts. Taking advantage of such unique expression pattern, we have developed a tamoxifen inducible Cre model, *Nr2f2-iCreERT2*, which specifically and efficiently targets interstitial cells in both male and female reproductive organs across embryonic and adult stages. This model offers a powerful tool for gene knockout studies specifically in the interstitial compartment, without affecting the supporting or germ cells. Additionally, it enables interstitial cell lineage tracing, facilitating the assessment of the interstitial’s contribution to non-interstitial cell types during development and differentiation.

## Introduction

The gonadal interstitium, despite its importance as the connective tissue that hold the organ together, is understudied compared to the supporting cell lineage (Sertoli and granulosa cells) and the germline. This is evident in the extensive mouse Cre models available to target these cell populations, including *Sf1-Cre, Sox9-Cre, Foxl2-Cre, Amh-Cre, Stra8-Cre,* and *Ddx4-Cre*, among others [1–6]. These models provided invaluable tools for exploring the biology of cell types critical to reproduction [7–14]. The interstitium of testis and ovary is essential to produce sex steroids that regulate spermatogenesis and folliculogenesis respectively [15–18]. Beyond hormone production, the interstitium provides structural integrity, vascular support, and facilitates paracrine signaling, enabling crosstalk with other gonadal cell types to sustain a functional reproductive system. Dysregulation of this compartment is implicated in several reproductive disorders, including hypogonadism, polycystic ovary syndrome (PCOS) and infertility [19–23]. Despite its critical functions, the study of interstitial cells has been limited by the lack of precise genetic tools.

Various Cre models have been used to target the gonadal interstitium under the control of key genes, including *Wt1*, *Sf1*, *Wnt5a* and *Tcf21*. *Wt1-CreERT2* is a tamoxifen-inducible Cre model that targets all somatic cells in the gonad, including interstitial and supporting cells, therefore lacking specificity [16, 24–27]. *Sf1-Cre* targets supporting and fetal Leydig cells, and a few (but not all) interstitial cells [1, 28]. *Tcf21^mCrem^*contributes broadly to all major somatic cells in the fetal testis, including Sertoli, Leydig and interstitial cells [29]. Lastly the *Wnt5a* model, which utilizes a Tet-On doxycycline-inducible system, might be the most specific for interstitial cells in the gonads [30]. However, its functionality depends on the presence of an additional transgene (TetO:Cre), requiring extra mouse breeding steps. Given these limitations, the development of a mouse model that specifically and efficiently targets interstitial cells is critical to advancing our understanding of the interstitial contributions to both normal physiology and pathological conditions in the gonad.

The orphan nuclear receptor *Nr2f2,* better known as COUP-TFII (Chicken Ovalbumin Upstream Promoter Transcription Factor II), functions as a transcription factor regulating the embryonic development of numerous organs [31]. *Nr2f2* has been shown to control several cellular processes including self-renewal, differentiation, and proliferation [32]. In the reproductive system, *Nr2f2* is expressed in ovary, testis, and uterus, where it plays essential roles in development and function [22, 33, 34]. In both male and female gonads, *Nr2f2* expression is specifically localized to the interstitial compartments, with little to no expression in supporting cells or germline cells [35, 36]. This unique expression pattern makes *Nr2f2* an ideal candidate for developing a Cre mouse line to enable tissue-specific and cell-specific genetic manipulations in interstitial cells of the ovary and testis. Moreover, *Nr2f2* is expressed in the stromal compartments of other reproductive structures, such as the Wolffian and Müllerian ducts and the adult uterus, broadening the utility of such a model for studying interstitial contributions across the reproductive system [37, 38].

## Materials and Methods

### Generation of *Nr2f2-T2A-iCreERT2* mice

The *Nr2f2-T2A-iCreERT2* locus was generated by replacing the genomic sequence at the stop codon in exon 3 with the improved iCre recombinase fused with the ERT2 tamoxifen-inducible domain and SV40 polyadenylation sequence (Figure 1A). The repair donor plasmid was generated in the pCR2.1 backbone by conventional molecular biology techniques and consists of a 760 bp 5’ homology (chr7:70,004,408-70,005,167 GRCm39), codon-improved iCreERT2 recombinase, SV40 polyadenylation signal, and a 912 bp 3’ homology arm (chr7:70,003,275-70,004,186GRCm39). In the final Nr2f2-T2A-iCreERT2 locus structure, 221 bp of endogenous sequence including the endogenous stop codon (chr7:70,004,187-70,004,407 GRCm39) was replaced by the T2A-iCreERT2-pA genetic payload (2225 bp).

**Figure 1:**
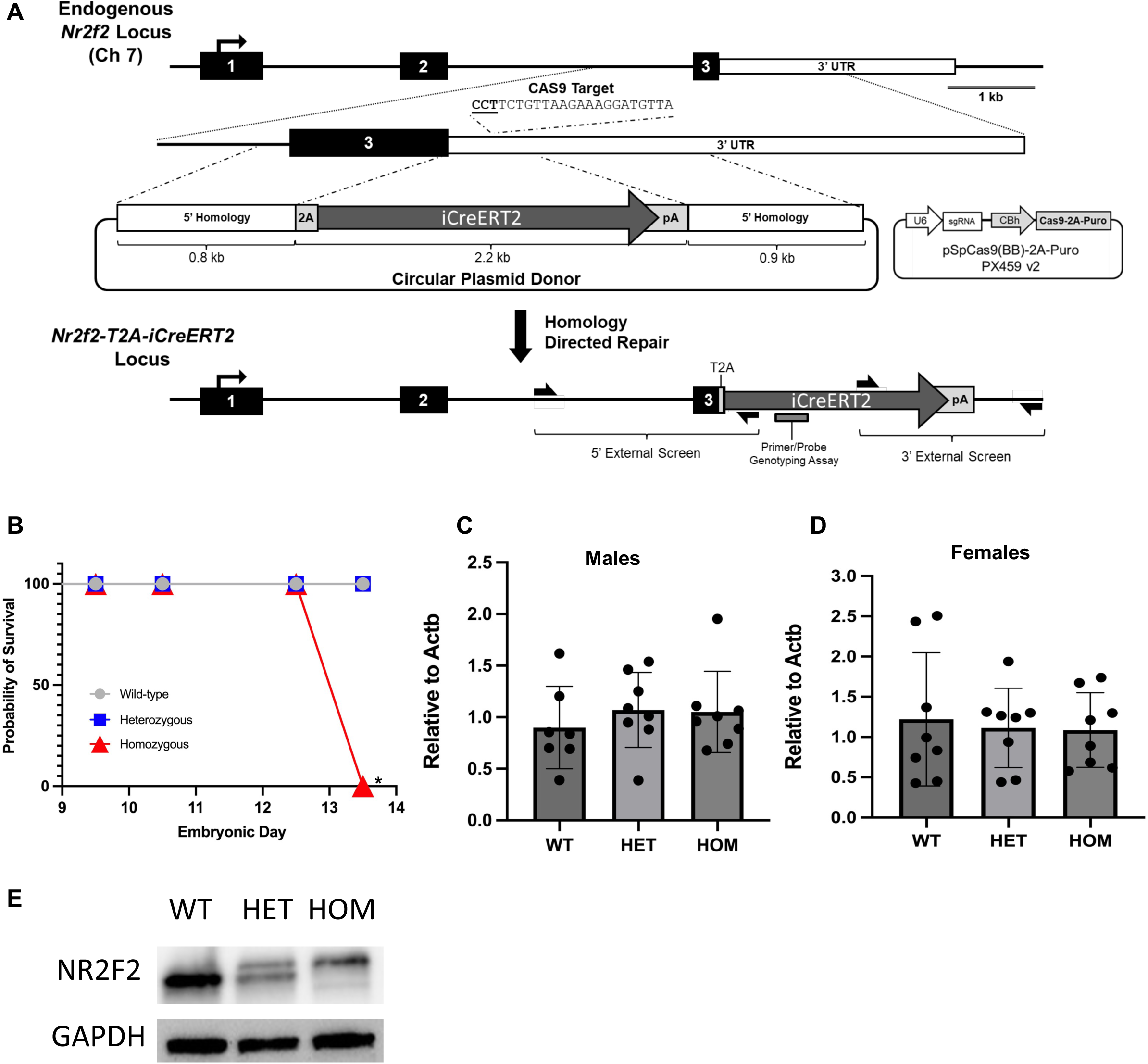
*Nr2f2-iCreERT2* knock-in generation and viability. (A) Schematic representation for generation of the *Nr2f2-iCreERT2* knock-in allele. T2A-iCreERT2-SV40 cassette was knocked into the endogenous *Nr2f2* allele just downstream of the stop codon using CRISPR technology. (B) Survival curve of *Nr2f2-iCreERT2* wildtype, heterozygous and homozygous embryos from E9.5 to E13.5. Mantel-Cox test; * p<0.05. (C & D) Quantitative RT-PCR quantification of *Nr2f2* mRNA expression, in male or female E12.5 gonad and mesonephros, relative to the housekeeping gene *Actb*. (D) Western blot detection of NR2F2 and GAPDH protein in the heads of the three different genotypes at E12.5. A representative of three biological replicates is shown. WT = wildtype, HET = one targeted allele, HOM = two targeted alleles.

Gene targeting was performed in B6129F1 embryonic stem cells (G4; 129S6/SvEvTac x C57BL/6Ncr). Embryonic stem cells were transfected with a 6:1 molar ratio of donor plasmid and Cas9-Puro/sgRNA (TAACATCCTTTCTTAACAGAAGG<u>NGG[PAM]</u>) delivery plasmid (pSpCas9(BB)-2A-Puro (PX459) V2.0), a gift from Feng Zhang (Addgene plasmid # 62988, PMC3969860).

After transfection, cells were selected using puromycin (0.9 µg/mL) followed by standard clonal expansion/screening. Clones were screened with 5’ and 3’ screens external to the homology arms and genetic payload/WT zygosity screen to identify Nr2f2-T2A-iCreERT2/WT clones. Screening primers: Nr2f2-Cre 5’Scr (Fwd: GTTTGCATGGCTGACCTTCAAT, Rev: CCACCTCTGATGAAGTCAGGAAG), Nr2f2-Cre 3’Scr (Fwd: CGAGTCCTGGACAAGATCACAG, Rev: CAAAGACTCGACCAAACAGTTCA), and *Nr2f2* WT Scr (Fwd: GTGGAAAGCTTGCAGGAAAAGT, Rev: ACAACTAGAGGTACATAGACACAGG). PCR amplicons from targeted clones were fully sequenced to confirm proper insertion of the T2A-iCreERT2-pA genetic payload. For the chimeric founder generation, targeted ES cells were microinjected into E3.5 albino B6J blastocysts (Jackson Laboratory stock #000058) isolated from naturally mated females. Injected blastocysts were then non-surgically transferred to pseudo-pregnant recipient SWISS mice. Germline male chimera founders were mated to C57BL/6J females to establish the *Nr2f2-iCreERT2* mouse line. The *Nr2f2-iCreERT2* allele was re-sequenced in the F1 chimera offspring. The line was maintained by backcrossing to C57BL/6J wildtype mice. The mouse colony was subsequently genotyped at Transnetyx using primer/probe assays; *Nr2f2* WT (Fwd: AGAGAGAAAGAGAGAGACTGCCAAA, Rev: TTCTGCCCAACACAGGAGTT, Probe: TTTAATGCATTCTGTAAAAGTTC) and iCre (Fwd: TCCTGGGCATTGCCTACAAC, Rev: CTTCACTCTGATTCTGGCAATTTCG, Probe: ACCCTGCTGCGCATTG).

### Effectiveness of CRE recombinase activity

Adult males heterozygous for the *Nr2f2-T2A-iCreERT2* allele were mated to *B6.Cg-Gt(ROSA)26Sor^tm9(CAG-tdTomato)Hze^* (or Rosa-tdTomato JAX #007909) females. The presence of a copulatory plug the following morning indicated gestational timing as embryonic day 0.5 (E0.5). Pregnant ROSA-tdTomato females were dosed daily from E11.5 to E13.5 intraperitoneally (I.P.) with 1.0mg/10g of body weight of tamoxifen (T-5648, Sigma-Aldrich) in corn oil, or corn oil alone as the control (MilliporeSigma). Embryos were collected at E14.5 and assessed for fluorescent expression using a Leica M165FC fluorescence dissecting microscope, Leica DFC310FC camera, and Leica Application Suite X (LAS X) software. Tissues were fixed in 4% paraformaldehyde (Electron Microscopy Sciences) for one hour at room temperature followed by multiple washes in 1x PBS buffer and stored at 4°C until processed. For inducing Cre activity in adult mice, 2 months old mice positive for both *Nr2f2-T2A-iCreERT2* and Rosa-tdTomato alleles were injected I.P. with 1.0mg/10g of body weight of tamoxifen for three consecutive days. Tissues were collected 48 hours after final tamoxifen injection, and fixed overnight in 10% neutral buffered formalin (MilliporeSigma) at 4°C and washed with 1x PBS multiple times the following day. All animal procedures were approved by the National Institute of Environmental Health Sciences (NIEHS) Animal Care and Use Committee in compliance with a NIEHS-approved animal study proposal (#2010-0016).

### Immunofluorescence

Embryonic tissues were processed in a Leica TP1020 tissue processor and embedded in paraffin blocks. Tissue sections were cut at six microns using a Leica RM2245 microtome and placed on positively charged glass slides. Sections were deparaffinized and rehydrated through ethanol series. Adult tissues were embedded in Tissue-Tek O.C.T. compound (Sakura) and sectioned at ten microns on a Leica CM3050S cryostat. Adult and embryonic sections were treated with a low pH citrate buffer (Vector Laboratories) to unmask protein epitopes. Primary antibodies, including Mouse anti-NR2F2 (R&D Systems, PP-H7147-00, 1:300), rabbit anti-dsRed (Takara, 632496, 1:1000), goat anti-AMH (Santa Cruz, sc-6886, 1:500), goat anti-CYP17A1 (Santa Cruz, sc-46081, 1:200) and goat anti-FOXL2 (Novus Biologicals, NB100-1277, 1:300), were incubated overnight at 4°C. Samples were washed in 1X PBS and incubated with fluorescent tagged secondary antibodies: donkey anti-rabbit 568 (Invitrogen, A10042, 1:500), donkey anti-rabbit 647 (Invitrogen, A31573, 1:500), donkey anti-goat 568 (Invitrogen, A11057, 1:500), donkey anti-goat 647 (Invitrogen, A21447, 1:500), donkey anti-mouse 568 (Invitrogen, A10037, 1:500) and donkey anti-mouse 647 (Invitrogen, A31571, 1:500), for one hour at room temperature. Slides were incubated with nuclear counterstain DAPI (MilliporeSigma) for five minutes at room temperature, coverslips applied with ProLong Diamond Antifade Mountant (Invitrogen), and images were captured using a Leica DMI4000 confocal microscope system running LAS X software.

### Quantitative RT-PCR analyses

Embryonic gonads with their mesonephros attached from time mated *Nr2f2*^iCreERT2/+^ males and females were collected from wildtype, heterozygous, and homozygous progeny aged E12.5. Total RNA from four biological replicates for each sex was isolated using an Arcturus Pico Pure kit (Applied Biosystems). cDNA was generated with SuperScript II Reverse Transcriptase (Invitrogen) according to manufacturer’s instructions. Expression levels of endogenous *Nr2f2* was determined using TaqMan (Thermo Fisher Scientific) probe Mm00772789_m1 in a Bio-Rad CFX96 Real-Time System. Data were analyzed using the ΔΔCT calculation normalized to ß-Actin (Mm04394036).

### Western blot for NR2F2 proteins

Protein was isolated from three biological replicates of each genotype of 12.5-day old male embryos using Pierce RIPA extraction buffer (Thermo Fisher Scientific) and homogenized for one minute with a handheld motorized pestle. Protein concentration was determined by BCA assay and read on a CLARIO Star Plus plate reader (BMG LabTech). Approximately 20µg of protein was loaded per lane of a Bio-Rad 10% TGX precast gel, run for an hour, and transferred to PVDF membrane using an iBLOT system (Invitrogen). Non-specific proteins were blocked in 5% dry milk overnight at 4 °C. The blot was then probed for endogenous NR2F2 using the same antibody listed above at 1:1000 dilution. HRP-conjugated secondary antibody was detected with a PICO Plus Chemiluminescent substrate (Thermo Fisher Scientific) kit following manufacturer’s recommendations and images captured on a Bio-Rad ChemiDoc MP. The blot was then stripped for 30 minutes at 37 °C in restore buffer (Thermo Fisher Scientific) prior to being probed with HRP-conjugated anti-GAPDH (Cell Signaling) as a loading control.

### Statistical analysis

Statistical analysis of embryonic survival of *Nr2f2*^icre/+^ progeny was determined using GraphPad Prism version 9.5.1 Mantel-Cox test. The RNA quantification data was also analyzed in GraphPad Prism. Any outliers were identified using ROUT (Q = 1%) prior to running an ordinary one-way ANOVA. One outlier was found and removed from the wildtype male samples. No statistical significance was found between the genotypes in their respective sex.

## Results

### Generation of the *Nr2f2-iCreERT2* mouse model

The *Nr2f2-T2A-iCreERT2* locus was generated by inserting a codon-improved *iCre* recombinase fused to the ERT2 tamoxifen-inducible domain into the 3’end of the stop codon in exon 3, to avoid creating a knockout allele (Figure 1A). The global *Nr2f2* knockout in mice is embryonically lethal at E10-E10.5 [39], we therefore evaluated if the incorporation of the CreERT2 cassette in the 3’ end of the *Nr2f2* gene might have caused any deleterious effects. Crosses between *Nr2f2-iCreERT2* heterozygous mice produced no homozygous offspring at birth, suggesting an embryonic lethality phenotype. We examined the survival rates of embryo at E9.5 and E10.5 (Figure 1B) and found that all embryos were viable at these timepoints. We then collected the embryos at later developmental stages and found that all homozygous embryos died between E12.5 and E13.5, suggesting a delayed mortality and potential residual expression of *Nr2f2* from the *Nr2f2-iCreERT2* locus (Fig. 1B).

To quantify the levels of *Nr2f2* expression across genotypes, we performed qRT-PCR for *Nr2f2* mRNA on male (Figure 1C) and female (Figure 1D) gonad-mesonephros samples, which are known to express *Nr2f2* [26, 35, 36]. No significant differences in *Nr2f2* mRNA expression were observed between the wildtype, heterozygous and homozygous samples. However, western blot analysis of E12.5 male brain tissue samples revealed the presence of a larger NR2F2 protein isoform in mice carrying the *Nr2f2-iCreERT2* allele (Figure 1E). This is likely to be a partial fusion protein as the T2A peptide has been reported to fail to cleave in other mouse models [40, 41].

### Detection of Nr2f2-iCreERT2 activity in embryos

To evaluate the activity and specificity of Nr2f2-iCreERT2, we injected tamoxifen or vehicle solution to pregnant females that carried control (*Nr2f2^+/+^; Rosa^tdTomato/+^*) and Cre-positive (*Nr2f2^+/iCreERT2^; Rosa^tdTomato/+^*) embryos at E11.5, E12.5, and E13.5. Embryos were examined for red tdTomato fluorescence, an indicator of Cre activity 24 hours after the last injection (E14.5) (Fig. 2A). As expected, no tdTomato expression was detected in any organs of the control embryos with or without tamoxifen (Fig. 2A, left). Conversely, *Nr2f2-iCreERT2* positive embryos exhibited variable levels of tdTomato expression only in the presence of tamoxifen (Fig. 2A, right). These results confirm the effectiveness and specificity of *Nr2f2-iCreERT2* in activating Cre-inducible reporters upon tamoxifen induction.

**Figure 2:**
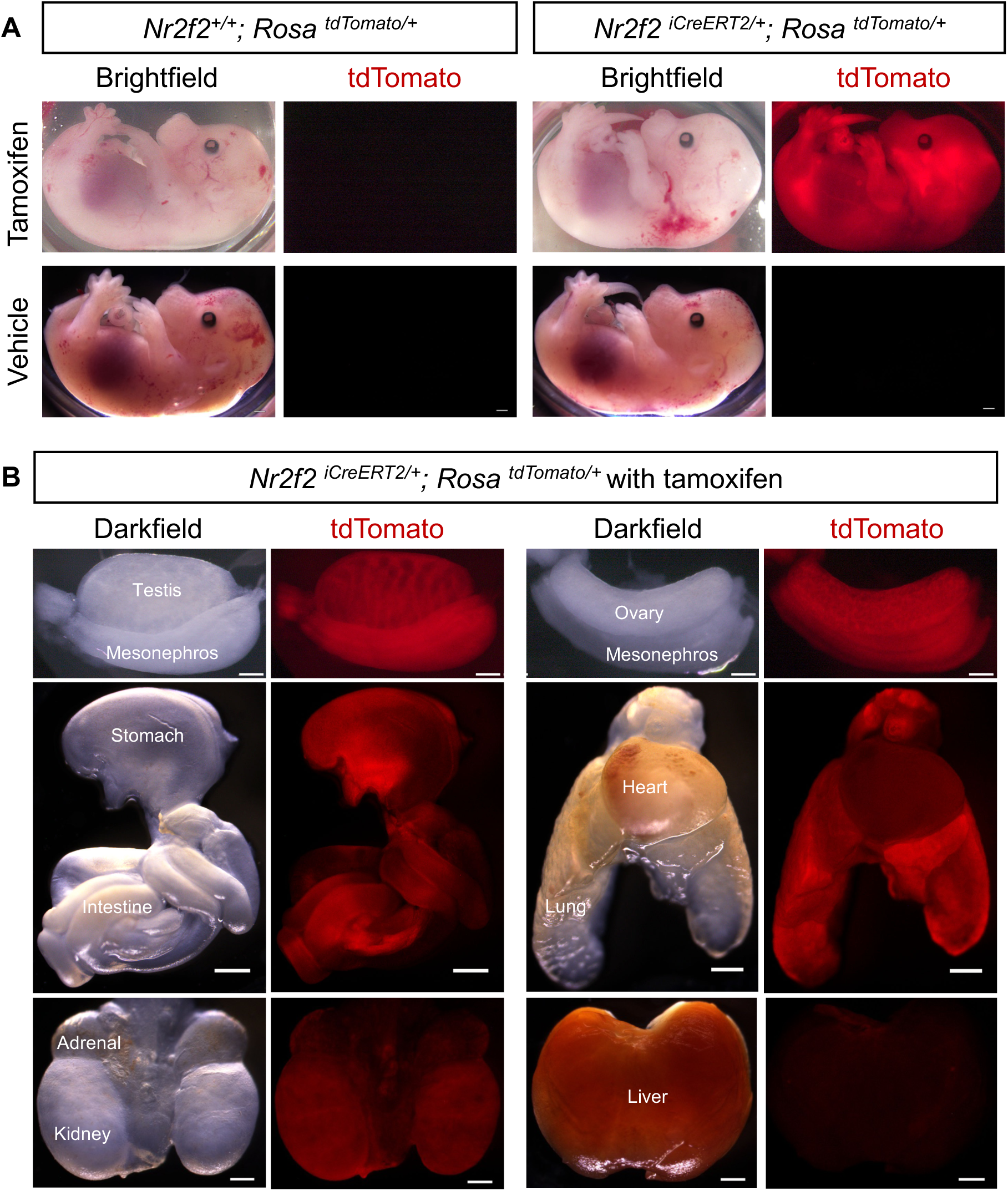
Activity of *Nr2f2-iCreERT2* in embryos. (A) Brightfield and tdTomato fluorescent expression in either control (*Nr2f2^+/+^; Rosa^tdTomato/+^*) or Cre-positive (*Nr2f2^iCreERT2/+^; Rosa^tdTomato/+^*) whole embryos treated with tamoxifen or vehicle solution. Scale bars 100 µm. (B) Darkfield and tdTomato fluorescent expression of various organs from an *Nr2f2-iCreERT2* positive mouse. Scale bars are 50 µm.

To determine whether tdTomato localization reflected native *Nr2f2* expression, we examined several organs from E14.5 *Nr2f2-iCreERT2* positive mouse embryos for red fluorescence. As expected, testis, ovaries and mesonephros were tdTomato positive (Fig. 2B), and tdTomato expression was also observed in stomach, intestine, kidneys and lungs, but not in adrenal, heart and liver (Fig. 2B). This pattern aligned with *Nr2f2 in-situ* hybridization data for E14.5 mouse embryos [42]. Next, we performed immunofluorescence of various cell markers on testes and ovaries of control and Cre-positive embryos at E14.5. As expected, no tdTomato expression was detected in control testes or ovaries exposed to tamoxifen (Fig. 3A and B). In the presence of tamoxifen, *Nr2f2-iCreERT2*-positive testes had cytoplasmic tdTomato in the interstitial compartment, but not in the testis cords where AMH+ Sertoli cells were located (Fig. 3A). In the interstitium, cytoplasmic tdTomato was co-expressed in nuclear NR2F2-positive cells in the interstitium, confirming the Cre specific expression under the *Nr2f2* promoter (Fig. 3A). Also, in the testis interstitium, some NR2F2-negative, CYP17A1-positive fetal Leydig cells were also tdTomato positive (Fig. 3A), suggesting that these Leydig cells initially expressed *Nr2f2* but subsequently downregulated it during differentiation. In *Nr2f2-iCreERT2*-positive ovaries, tdTomato was expressed in NR2F2-positive interstitial cells, but not in FOXL2-positive granulosa cells (Fig. 3B). These data confirmed the correct and specific *iCreERT2* activity under the *Nr2f2* promoter in response to tamoxifen, and its ability to target interstitial cells.

**Figure 3:**
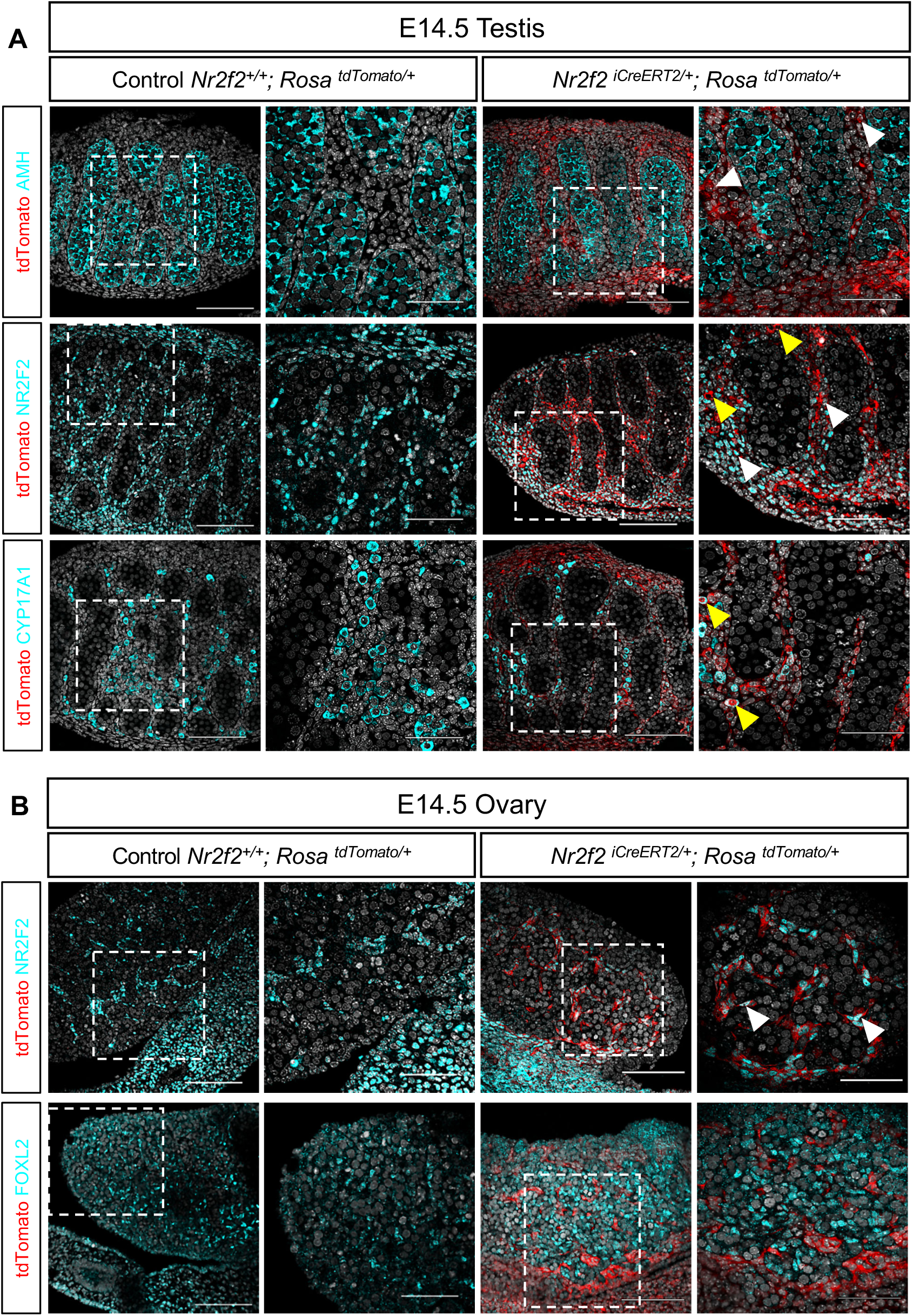
*Nr2f2-iCreERT2* activity in interstitial cells in the embryonic gonad. (A) immunofluorescence on testicular sections from either a control or *Nr2f2-iCreERT2* positive mouse probed with antibodies to tdTomato (red) and endogenous AMH (Sertoli cells), NR2F2 (interstitial cells) or CYP17A1 (fetal Leydig cells). (B) Immunofluorescence on ovarian sections from either a control or *Nr2f2-iCreERT2* positive mouse probed with antibodies to tdTomato (red) and endogenous NR2F2 (interstitial cells) or FOXL2 (pre-granulosa cells). Dashed white box indicates magnified area. Scale bars are 100 µm and 50 µm (Magnified area). White arrowheads indicate tdTomato positive interstitial cells. Yellow arrowheads indicate NR2F2 negative, CYP17A1 positive, tdTomato positive interstitial cells.

### Nr2f2-iCreERT2 activity in adult tissues

To determine whether *Nr2f2-iCreERT2* could also be used to target adult reproductive organs, we injected the control (*Nr2f2^+/+^; Rosa^tdTomato/+^*) and Cre-positive (*Nr2f2^+/CreERT2^; Rosa^tdTomato/+^*) adult mice (2 months old) with tamoxifen for three consecutive days to induce tdTomato reporter expression. Tissues were harvested 48 hours after the last injection. Positive tdTomato expression was detected in the male reproductive system, including testes, epididymis, seminal vesicles and the bladder; and in the ovaries and uteri of female mice (Fig. 4A). Positive tdTomato expression was also observed in the lung, kidney, adrenal glands and liver, but not in the heart or gallbladder (Fig. 4B). Next, we performed immunofluorescence of various cell markers on testes, ovaries, and uteri of control and Cre-positive adult animals (Fig. 5). As expected, control animals showed no tdTomato expression in the presence of tamoxifen. In the Cre-positive testes, tdTomato expression was observed in the interstitial space that surrounds the seminiferous tubules, but not inside the tubules (Fig. 5A). In the epididymis, expression patterns varied across regions: positive tdTomato expression was found in the interstitial space and in the muscle layer underneath the epithelium in the caput and epididymis. Positive tdTomato staining was also observed within the caput epithelium (Fig. 5B). In the adult ovary, tdTomato positive staining was observed in the stroma and theca cells (Fig. 5C) but was absent in granulosa cells and oocytes (Fig. 5D). In the uterus, tdTomato staining was present in the endometrial stroma, but not in the glandular epithelium. Notably, some uterine epithelial cells exhibited tdTomato expression, despite NR2F2 not being expressed in this cell type [38]. This suggests a mesenchymal origin for some of the uterine epithelial cells.

**Figure 4:**
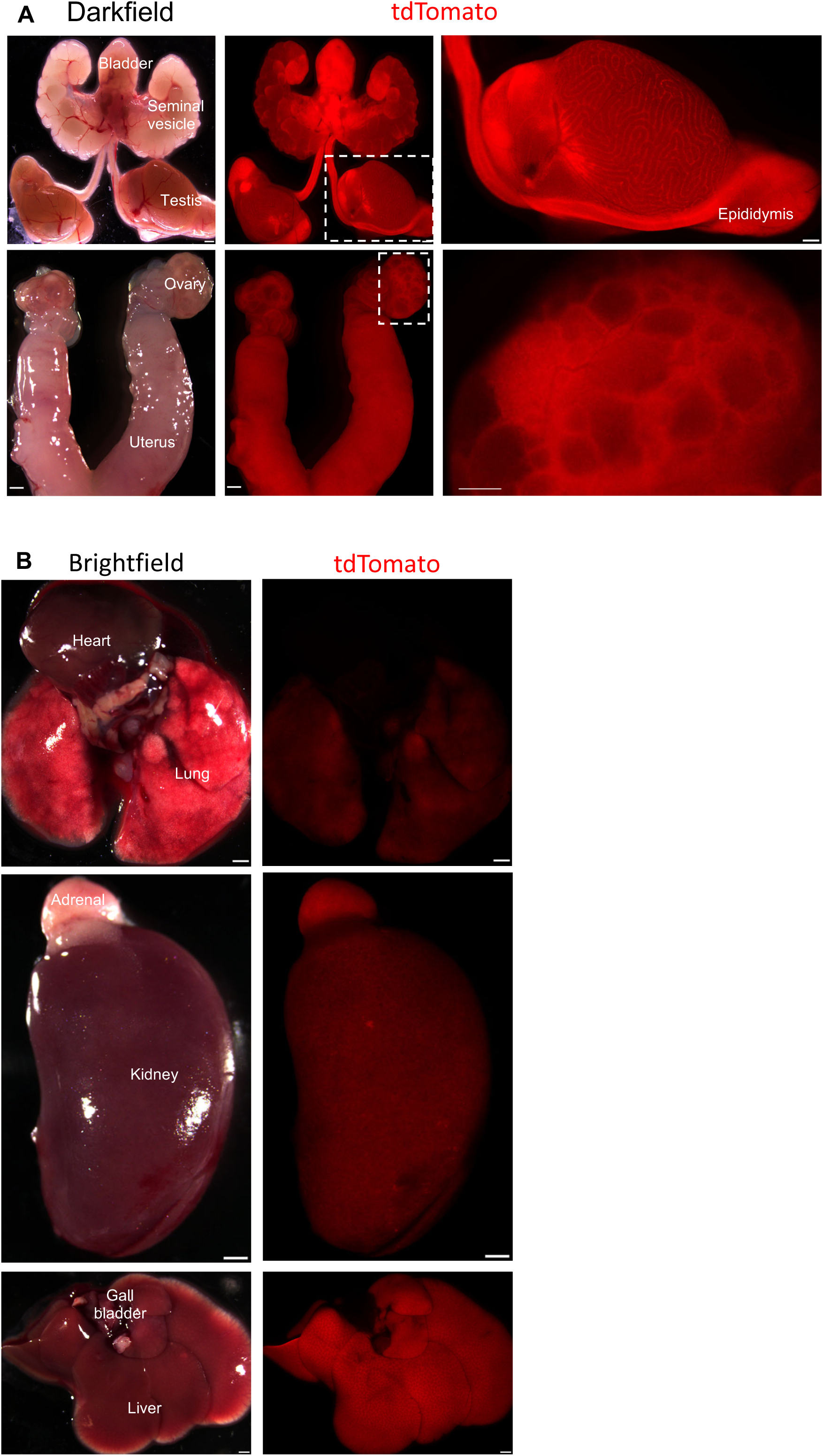
Activity of *Nr2f2-iCreERT2* in adult tissues. (A). tdTomato fluorescent expression in *Nr2f2-iCreERT2* male and female reproductive systems. (B) tdTomato fluorescent expression in *Nr2f2-iCreERT2* various organs. Scale bars are 100 µm. Dashed white box indicates magnified area.

**Figure 5:**
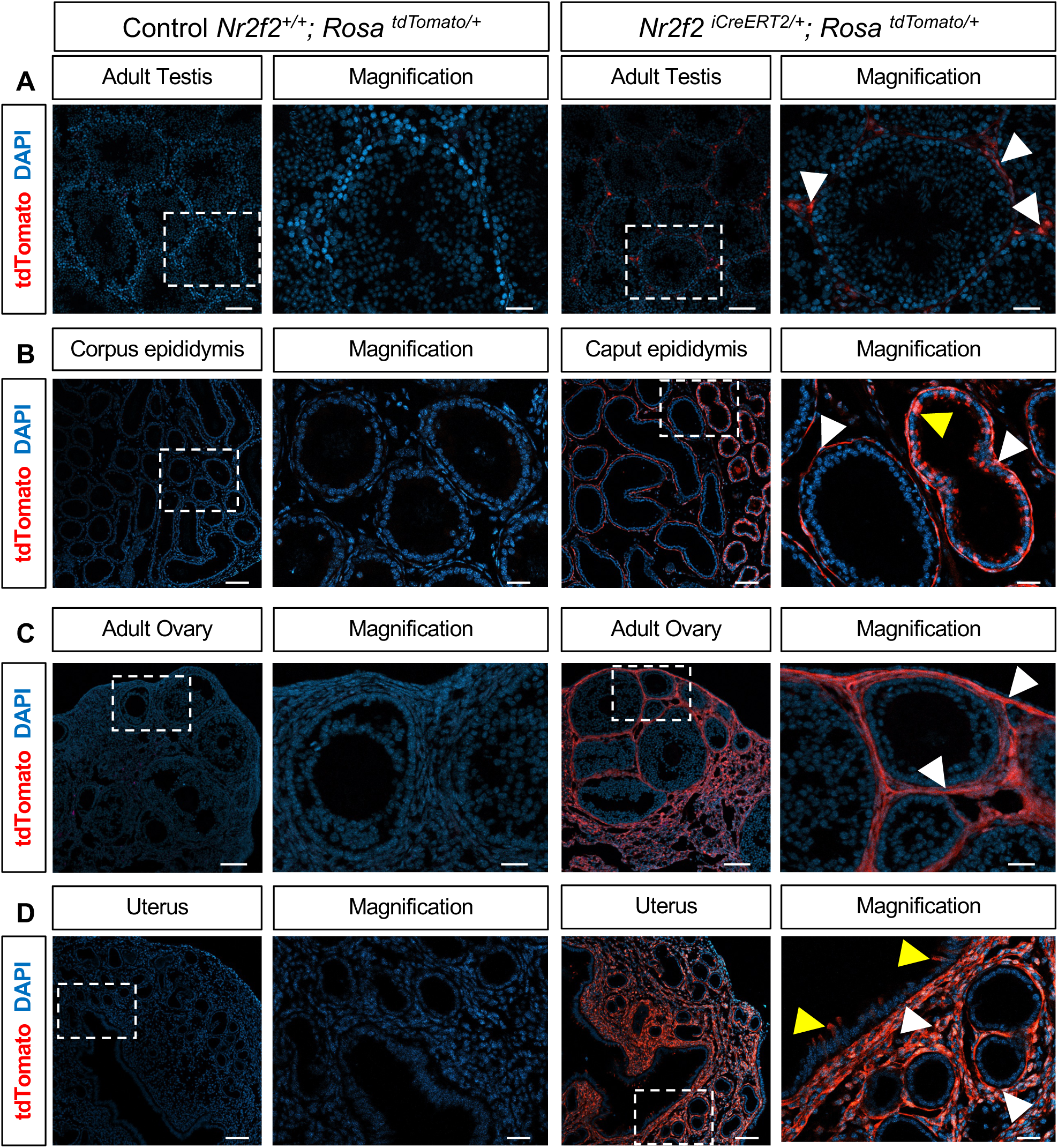
*Nr2f2-iCreERT2* activity in interstitial cells in the adult reproductive systems. Immunofluorescent detection of tdTomato (Red) in testicular (A), epididymal (B), ovarian (C) and uterine (D) sections from either wildtype or *Nr2f2-iCreERT2* positive mice. Dashed white box indicates magnified area. White arrowheads indicate tdTomato positive interstitial cells. Yellow arrowheads indicate, tdTomato positive epithelial cells. Scale bars are 100 µm and 33 µm (Magnified).

## Discussion

We successfully generated a new inducible Cre mouse line that can be used to specifically target the interstitial/stromal cells in the reproductive systems. Due to the apparent importance of the supporting gonadal cells and germline in reproduction, most of the Cre mouse models in gonadal research have focused on these two cell populations. However, this emphasis has often overshadowed the importance of interstitial cells, which are essential for fertility and reproduction [15–23, 43]. This highlights the need for specific mouse models to target these interstitial cells. One of the most commonly used inducible Cre models to target the interstitial cells is the *Wt1-CreERT2* [16, 24–27]. However, this model lacks specificity as it also targets the supporting cell lineage and displays haploinsufficiency, which could confound the observed phenotypes. In recent years, several groups attempted to generate reliable animal models to target the gonadal interstitium. Two inducible animals were commonly used to target the interstitial populations in the gonad, the tamoxifen inducible *Tcf21-Cre*, and the *Wnt5a Tet-on* system [29, 30]. Unfortunately, neither of them is commercially available. Other animal models can target only a subset of the interstitial cells, for example *Cyp17a1-Cre* targets Leydig cells whereas *Myh11-Cre* targets peritubular myoid cells [23, 44].

In this study, we generated a new model that specifically targets the gonadal interstitial cells using a tamoxifen inducible, enhanced version of Cre (iCreERT2) that were knocked into the 3’ UTR of the endogenous *Nr2f2* gene, a known gonadal interstitial cell marker [36, 45]. Our model targets the interstitial lineage, but not the supporting cell compartments. Moreover, this model is active in the interstitial lineages in both embryonic and adult male and female tissues, showcasing the versatility of the model. Although we reported the use of three consecutive tamoxifen injections, we also found a single dose to be sufficient to label a substantial number of interstitial cells in embryonic testis [26], confirming the efficiency of this model.

While this model predominantly labels interstitial cells, a few epithelial cells were also labeled in the reproductive tracts. Notably, these epithelial cells do not express *Nr2f2* [38]. This suggests that interstitial cells may contribute to the epithelial cell population during tissue homeostasis, potentially through a mesenchymal-to-epithelial transition that involves the downregulation of *Nr2f2*. A similar pattern was observed in fetal Leydig cells, where NR2F2-negative/CYP17A1-positive fetal Leydig cells were labelled with tdTomato, indicating that fetal Leydig cell progenitors initially express *Nr2f2*, and then downregulate its expression during fetal Leydig cell differentiation. These observations showcase the role of interstitial cells as progenitors and demonstrate how this model can be used to study cell differentiation.

Global knockout of *Nr2f2* resulted in lethality by E10-E10.5 due to hemorrhage and edema in the brain and heart [39]. Although our knock-in strategy was designed to insert the iCreERT2 cassette after the *Nr2f2* stop codon, without altering its function, the mice exhibited a higher molecular weight for NR2F2 protein compared to controls. This suggests the presence of an alternative splicing isoform or post-transcriptional modification, possibly a T2A retention. Despite this aberrant protein, the homozygous knock-in embryos did not die at E10, but rather between E12.5 and E13.5, suggesting that modified NR2F2 proteins retain a residual but less effective activity (a potential hypomorphic allele). Importantly, the heterozygous animals were normal and fertile, and our model exhibited no leakiness, as tdTomato expression was not detected in the absence of tamoxifen and it was confined to the expected expression sites of *Nr2f2*.

In conclusion, we have developed an effective inducible Cre mouse line capable of specifically targeting interstitial cells in both male and female reproductive organs at embryonic and adult stages. This model is ideal for knocking out genes specifically in the interstitial compartment, without affecting the supporting or the germline. Additionally, it can be used for interstitial cell lineage tracing to evaluate the contribution of the interstitium to other non-interstitial cell types during development and differentiation. Given that *Nr2f2* is expressed in multiple tissues during embryonic development, including the lung, the kidney, adrenal glands and the liver, this model holds great potential for applications beyond the reproductive field, benefiting the broader research community.

## Acknowledgments

We are grateful to the NIEHS Gene Editing and Mouse Model Core Facility for creating the mouse model and Comparative Medicine Branch for mouse colony husbandry. We also want to thank all the members of the Yao Laboratory for their continuous advice and encouragement. This work was supported by the Intramural Research Program (ES102965 to H.H.-C.Y. and ES102425 to MKR) of the NIH, National Institute of Environmental Health Sciences.

## Author contributions

H.Y., P.B. and M.R conceived the project and designed the experiments. P.B., M.E, A.G., G.S and T.H conducted experiments. M.E., P.B and H.Y. wrote the manuscript. All authors edited and revised the text.

## Data Availability

The authors confirm that the data supporting the findings of this study are available within the article and by request to the corresponding authors.

